# Onion thrips *Thrips tabaci* [Thysanoptera: Thripidae] reduces yields and THC in indoor grown cannabis

**DOI:** 10.1101/2020.05.11.088997

**Authors:** Frédéric McCune, Chad Murphy, James Eaves, Valérie Fournier

## Abstract

Cannabis (*Cannabis sativa* L. [Rosales: Cannabaceae]) is a newly legalized crop and requires deeper insights on its pest communities. In this preliminary study, we identified a thrips species affecting indoor grown cannabis in Canada and tested its impact on plant yield. We used three levels of initial infestation (zero, one, and five thrips) on individual plants grown in two growing mediums: normal substrate or substrate containing the biostimulant *Bacillus pumilus*, Meyer and Gottheil [Bacillales: Bacillaceae]. We found that the onion thrips, *Thrips tabaci* (Lindeman) [Thysanoptera: Thripidae] is proliferating in indoor grown cannabis. Furthermore, our results showed that fresh yields were higher for the plants that initially received zero thrips compared to those that initially received five thrips. Moreover, the biostimulant did not help reduce the impact of thrips. We highlight the importance for growers to carefully monitor thrips infestations in indoor grown cannabis. Finally, we emphasize the need for more research related to the impact of pests on cannabis yields and safe means of pest control for this strictly regulated crop.

## RÉSUMÉ

Le cannabis (*Cannabis sativa* L. [Rosales: Cannabaceae]) est une culture nouvellement légalisée et qui requiert des connaissances approfondies sur ses ravageurs. Dans cette étude préliminaire, nous avons identifié une espèce de thrips affectant le cannabis cultivé à l’intérieur au Canada et testé son impact sur le rendement des plants. Nous avons testé trois niveaux initiaux d’infestation (zéro, un, et cinq thrips) sur des plants individuels cultivés dans deux terreaux: un substrat normal ou un substrat contenant le biostimulant *Bacillus pumilus*, Meyer and Gottheil [Bacillales: Bacillaceae]. Nous avons observé que le thrips de l’oignon, *Thrips tabaci* (Lindeman) [Thysanoptera: Thripidae] prolifère dans le cannabis cultivé à l’intérieur. Nos résultats montrent que le rendement des plants de cannabis est plus élevé pour les plants n’ayant pas reçu de thrips comparativement aux plants sur lesquels cinq thrips ont initialement été inoculés. De plus, le biostimulant n’a pas permis de réduire l’impact des thrips. Nous mettons en lumière l’importance pour les producteurs de cannabis cultivé à l’intérieur de faire un suivi rigoureux de leurs populations de thrips. Finalement, nous soulignons les besoins importants en recherches concernant les ravageurs du cannabis, leurs impacts et le développement de méthodes de lutte dans cette culture hautement règlementée.

## INTRODUCTION

Cannabis (*Cannabis sativa* L. [Rosales: Cannabaceae]) was legalized for recreational purposes in October 2018 in Canada and is still under strict prohibition in most of the world. Thus, there is a severe lack of information regarding its growing practices (Eaves, Eaves, Morphy, & Murray, 2020; Wilson et al., 2019). This includes research related to the impact of pest species and the means of controlling them (Cranshaw et al., 2019). Under the Cannabis Regulations and the Pest Control Products Act, Health Canada only allows cannabis growers to use a limited number of pesticide products. Consequently, companies rely mostly on biological control, but these techniques are very costly, increase the risk of contaminating the final product with dead insect parts, and yield uneven results.

More than 300 arthropod species have been identified on hemp and cannabis (Cranshaw et al., 2019; McPartland, 1996a). On cannabis, the most predominant ones are sap-sucking arthropods such as aphids, whiteflies, leafhoppers, mealybugs and various mites (Lago & Stanford, 1989; McPartland, 1996a; Wilson et al., 2019). Recent reports of potential pests in cannabis include the marmorated stink bug (*Halyomorpha halys*) (Britt, Pagani, & Kuhar, 2019) and two aphid species (*Phorodon cannabis* and *Rhopalosiphum rufiabdominale*) (Lagos-Kutz, Potter, DiFonzo, Russell, & Hartman, 2018). Despite this, it is believed that very few insects can actually cause significant losses in commercial cannabis production (Dewey, 1913; McPartland, 1996a). In a recent survey, growers from California reported from zero to over 25% crop damage caused by arthropods (Wilson et al., 2019). Nonetheless, a large proportion of cannabis production occurs indoor or in greenhouses, which provide environments that are particularly favourable for pests. If fact, in Canada, Health Canada only started licensing outdoor area in October of 2019. Since most published studies have focused on outdoor production, our current estimates of pest-risk posed to cannabis producers may greatly underestimate the actual risk.

Thrips have been shown to be a major pest for many crops, most notably in greenhouses (Stuart, Gao, & Lei, 2011) and can inflict both direct and indirect damage (Hao, Shipp, Wang, Papadopoulos, & Binns, 2002; Pereira et al., 2017). Damage resulting from sucking or ovipositing in the marketable plant parts, like fruits, correspond to direct damage (Shipp, Hao, Papadopoulos, & Binns, 1998), while damage caused on non-marketable plant parts, like leaves, are considered indirect damage (Diaz-Montano, Fuchs, Nault, Fail, & Shelton, 2011; German, Ullman, & Moyer, 1992). Thrips are found in indoor cannabis facilities (McPartland, 1996a) but we are not aware of any studies investigating their impact on cannabis yields. Nevertheless, Cranshaw et al. (2019) reports that thrips are common pests in hemp farms. For instance, Onion thrips, *Thrips tabaci* (Lindeman) [Thysanoptera: Thripidae] have been frequently found on hemp in Colorado and can cause important foliage damage on indoor grown plants (Cranshaw et al., 2019). Western flower thrips, *Frankliniella occidentalis* (Pergande), tobacco thrips, *Frankliniella fusca* (Hinds) and greenhouse thrips, *Heliothrips haemorrhoidalis* (Bouché) have also been found in hemp farms (Cranshaw et al., 2019; Lago & Stanford, 1989; McPartland, 1996a).

Biostimulants are biological products that improve the productivity of plants. These products are often a mixture of compounds derived from various organisms, such as bacteria, fungi, algae, higher plants or animals, and frequently possess unexplained modes of action (Calvo, Nelson, & Kloepper, 2014; Conant, Walsh, Walsh, Bell, & Wallenstein, 2017; Yakhin, Lubyanov, Yakhin, & Brown, 2017). Specifically, the bacterium *Bacillus pumilus*, Meyer and Gottheil, [Bacillales: Bacillaceae] is known for its growth promoting (de-Bashan, Hernandez, Bashan, & Maier, 2010; Gutiérrez-Mañero et al., 2001; Probanza, Lucas, Acero, & Gutierrez Mañero, 1996) and antifungal (Pérez-García, Romero, & de Vicente, 2011) properties. Furthermore, *B. pumilu*s successfully suppressed larvae of *Scirpophaga incertulas* and *Bruchus dentipes* in laboratory conditions (Rishad, Rebello, Shabanamol, & Jisha, 2017; Tozlu, Dadasoglu, Kotan, & Tozlu, 2011). These results are likely explained by its high production of chitinase, an enzyme that can degrade the chitin containing cell walls of insects and thus induce death (Rishad et al., 2017). Chitinase has shown insecticidal properties against weevils (Laribi-Habchi, 2014) and aphids (Kim & Je, 2010). Growing mediums enhanced with entomopathogenic bacteria represents a promising avenue toward pest control and reduced use of pesticides. When added to a growing medium, *B. pumilus* reduces the infestation level of fungus gnats (Diptera) in greenhouses, but shows inconclusive results for the western flower thrips (Gravel & Naasz, 2019).

Hence, the objectives of this preliminary study were to identify the thrips species affecting indoor cannabis production in Ontario, to determine the potential yield and quality losses associated with their infestations, and, finally, to evaluate the impact of adding the biostimulant *B. pumilus* to the growing substrate on the impacts of thrips.

## MATERIALS AND METHODS

### Experiment

The experiment was conducted in the autumn of 2019 in the commercial cannabis production facility of GreenSeal Cannabis Company located in Stratford, Ontario, Canada. We used 60 clones (approximately two weeks old) of cannabis (*C. sativa* var. Green Crack) to test the impact of three initial levels of thrips infestation (zero, one or five thrips) and two growing substrates (normal or biostimulant). Ten cannabis clones, acting as ten replicas, were randomly assigned to each combination of infestation level and growing substrate. All clones were planted in seven inches square pots (4L) using one of the two types of substrate. The first (“normal substrate”) was a fibrous, peat-moss substrate with perlite (Pro-Mix HP Mycorrhizae, Premier Tech). The second (“biostimulant substrate”) was the normal substrate with the addition of the biostimulant *Bacillus pumilus* (strain GHA180) (Pro-Mix HP Biostimulant + Mycorrhizae, Premier Tech). Plants were propagated in a quarantine room and we visually inspected them for predatory mite or thrips. As an additional precaution, we carefully used a spray-bottle and cloth to wipe each individual leaf to ensure no arthropods were on them.

As thrips can reproduce asexually (Morison, 1957; Stuart et al., 2011) inoculating a single immature thrips can lead to an significant population overtime. It is thus not possible to control for their final infestation levels. Even though we do not think that controlling the initial infestation levels will result in consistent levels of infestation at the end of the experiment, we consider that it provides a valuable insight about the impact of a pest (Torres-Vila, Rodríguez-Molina, & Lacasa-Plasencia, 2003). In this way, we inoculated the plants with zero, one and five thrips to represent respectively control, low and high levels of infestation (Hao et al., 2002). Thrips used for the experiment were collected directly from the production area of the facility with entomological mouth aspirators. We targeted what we believed to be late instar larvae. Thrips were carefully inoculated on each plant with fine brushes. All plants, including the controls with no inoculated thrips, were then covered with Nitex (150 µm mesh) bags that were supported by stainless-steel frames and tightly secured around the pots by elastics bands. The fine meshes of the Nitex bags help prevent thrips from escaping and non-experimental pests or predatory mites from entering. Nonetheless, since this experiment was conducted in a production facility, rather than inside a university lab (It is still very difficult to receive a cannabis research license in Canada.), we did expect some cross contamination and thus recorded leaf damage for all plants. Two drippers were threaded under the elastic bands for irrigation purpose. We considered monitoring the thrips levels over the course of the experiment but decided the risk of allow arthropodes to enter or escape was too high.

The plants were grown in the facility’s quarantine room, where all plants were placed on two-levels shelves. Five plants of each treatment were placed on each level of the shelves following a completely randomized design, so that 30 plants were located on the top level and 30 plants on the bottom level. Plants were placed in two rows of 15 plants on each level. Shelves were equipped with broad-spectrum LED lighting (Voltserver High-Intensity Lighting Platform). Light intensity was gradually increased each week from 25% intensity (average of around 500 PPFD) to a maximum of 50% intensity (average of around 1,080 PPFD). Plants were then kept under commercial cultivation conditions (day/night temperature of 25°C/21°C +/- 2, 12 hours of daylight, 50% +/- 5 RH, and CO2 at ambient levels). Using a short vegetative period combined with these environmental conditions are typical for growers who follow a “sea of green” growing strategy, as GreenSeal Cannabis does. The exception is CO2 concentration levels, which were maintained at ambient levels for the experiment. Plants were watered by drip irrigation about every two days with approximately 1 L of water/plant.

All 60 plants were grown in the cages under aforementioned conditions for eight weeks. At the end of the eight-week period, the plants had reached the end of the flower stage. The fresh inflorescences were then harvested following normal commercial methods and weighed for each plant (Pennsylvania 7600 Series Bench Scale 4536 g x 0.5 g). In order to measure total THC levels, we took flower samples from three plants from each treatment with zero or five thrips. The samples for each treatment were then blended together and analyzed using HPLC analysis conducted by A&L Canada Laboratories Inc. located in London, Ontario. We therefore obtained a single THC measure for each control and high infestation treatment.

Multiple rows of plants in the two production rooms of the cannabis facility were checked to collect and identify all thrips species occurring in the facility. All collected specimens were observed under a stereomicroscope and appeared to be similar. Multiple thrips specimens at both adult and larval stages were collected and sent to the expert insects taxonomist from the Laboratoire d’expertise et de diagnostic en phytoprotection (LEDP) of the Ministère de l’Agriculture, des Pêcheries et de l’Alimentation du Québec for identification (Palmer, Mound, Du Heaume, & Betts, 1989), who has stored the vouchers. No other pests than thrips were observed in the facility.

### Statistical analyses

Fresh yield data were analyzed with R (R Core Team, 2019). We used generalized least squares fitted linear models and linear mixed-effects models (package “nlme” (Pinheiro, Bates, DebRoy, Sarkar, & R Core Team, 2019)). Fresh yield was used as the response variable, while both the number of thrips initially inoculated and the substrate type were explanatory variables. All models included all interactions between our explanatory variables. We first computed generalized least squares fitted linear models and then compared these to linear mixed-effects models in which the shelves’ level (upper or lower) was treated as a random effect. This allows us to take into account a potential difference in temperature or growing conditions between levels. We thereafter compared both types of models based on their Akaike information criterion (AIC). The more complex linear mixed-effects models including the shelves’ level had a higher AIC than the more simple models, indicating less accuracy of the model. We therefore only present results from the generalized least squares fitted linear models in the result section. We used an ANOVA to test the effect of both the number of inoculated thrips and the type of substrate on the fresh yield. We changed the reference level and fitted models once more to evaluate the effect of the multiple levels of our explanatory variables that were identified as significant in the ANOVA.

## RESULTS

All thrips specimens collected in the GreenSeal Cannabis Company’s facility were identified as being onion thrips, *Thrips tabaci* (Lindeman). The final fresh yield of our individual plants varied between 36.81 g and 195.84 g. Three plants died at the start of the experiment from transplant shock. We did not have replacement plants. They were respectively under treatments zero thrips-biostimulant, one thrips-normal, and five thrips-normal. Five others grew far more slowly than the average plant, which may indicate they were somehow stunted. Those were respectively under treatments zero thrips-biostimulant, zero thrips-normal (two plants), one thrips-biostimulant, and one thrips-normal. The plants that died or had a reduced growth were thus represented in almost all treatments, but slightly more in controls and low infestation treatments. Nonetheless, a boxplot revealed that only three of these observations were actual outliers. We removed the dead plants from our dataset and performed all analyses with and without the stunted plants. We obtained very similar results both times and thus decided to include the stunted plants in all analyses. We observed thrips on most plants at the end of the experiment, even on many control plants. However, we are not very concerned about this contamination since the relative amount of damage is representative of our desired levels of infestation. For example, control plants had very little damage compared to what was observed on the treatment plants. For the plants inoculated with five thrips, the total THC level was of 17.6% for the normal substrate and 17.77% for the biostimulant substrate. For the zero thrips treatment, the total THC level was 19.34% for the normal substrate and 19.62% for the biostimulant substrate. We limited our THC measurements to those four samples and thus cannot provide statistics here. Those THC levels nevertheless indicate a possible reduction in THC when plants are under high infestation of thrips.

The number of thrips had a significant effect on the final yield of the cannabis plants (ANOVA F(2, 51) = 7.1062, P = 0,0019; Fig. 1). Specifically, the yields were lower for plants that were initially inoculated with five thrips compare to the plants that had no thrips (t(51) = −2.569502, P = 0.0132). Average yield was 30.68% higher for plants that were not inoculated with thrips (130.15 g per plant compared to 90.22 g). This result highlights the relevance of keeping stunted plants in the analysis. Indeed, as the prevalence of stunted plants was slightly higher in the control and low infestation treatment, the effect of including these low yielding plants in the models was to reduce the potential negative impact of thrips. As we found an opposite trend, it therefore reinforces the hypothesis that thrips negatively impact yields. The effect of the substrate type was not significant (ANOVA F(1, 51 = 3.5769, P = 0.0643; Fig. 2) while the interaction between the infestation level and the substrate type (ANOVA F(2, 51 = 0.0506, P = 0.9507) had no effect on the final yields.

**Fig. 1.**
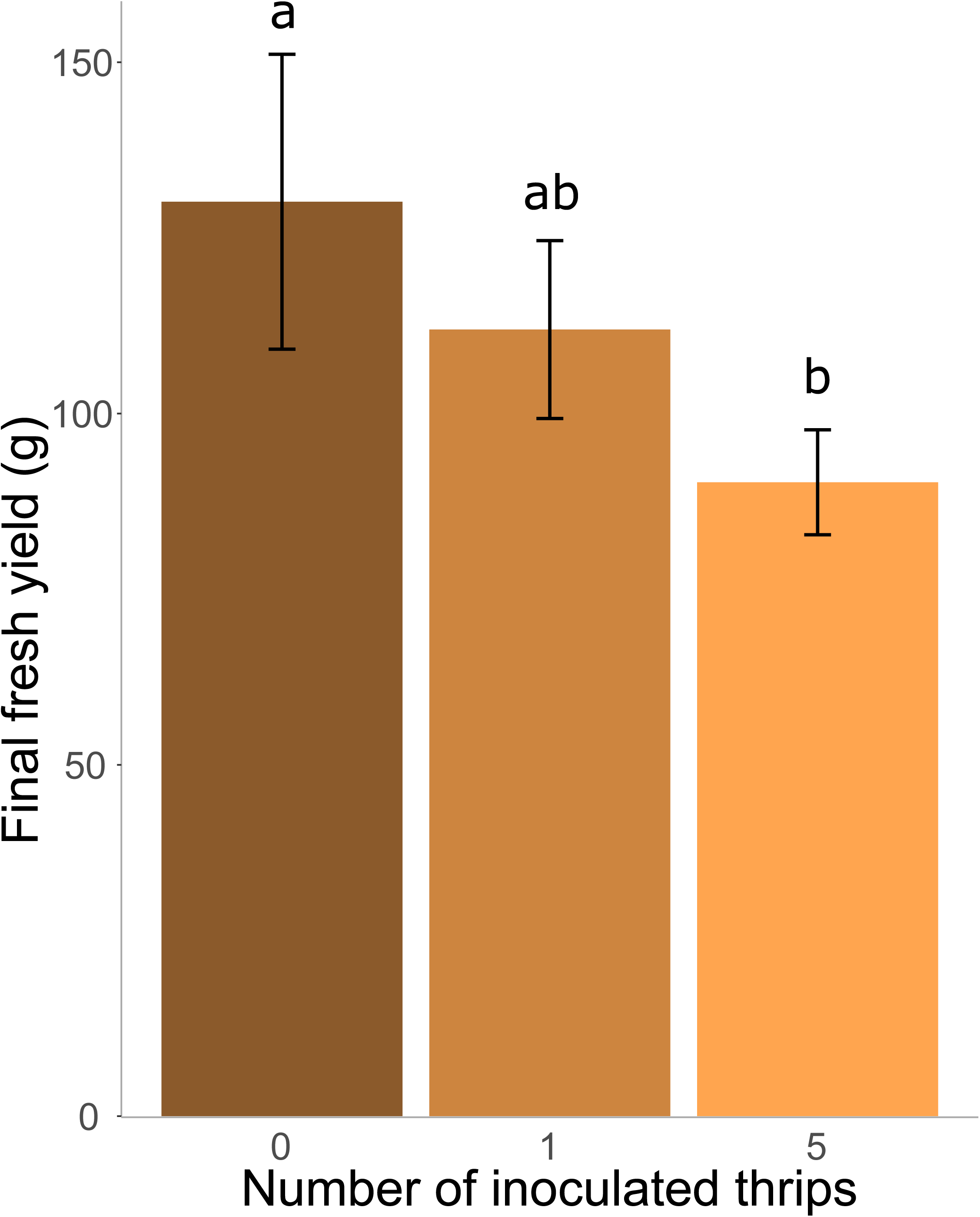
Mean final fresh yields of the cannabis plants after eight weeks according to the number of inoculated onion thrips (*Thrips tabaci*). Error bars represent 95% confidence intervals.

**Fig. 2.**
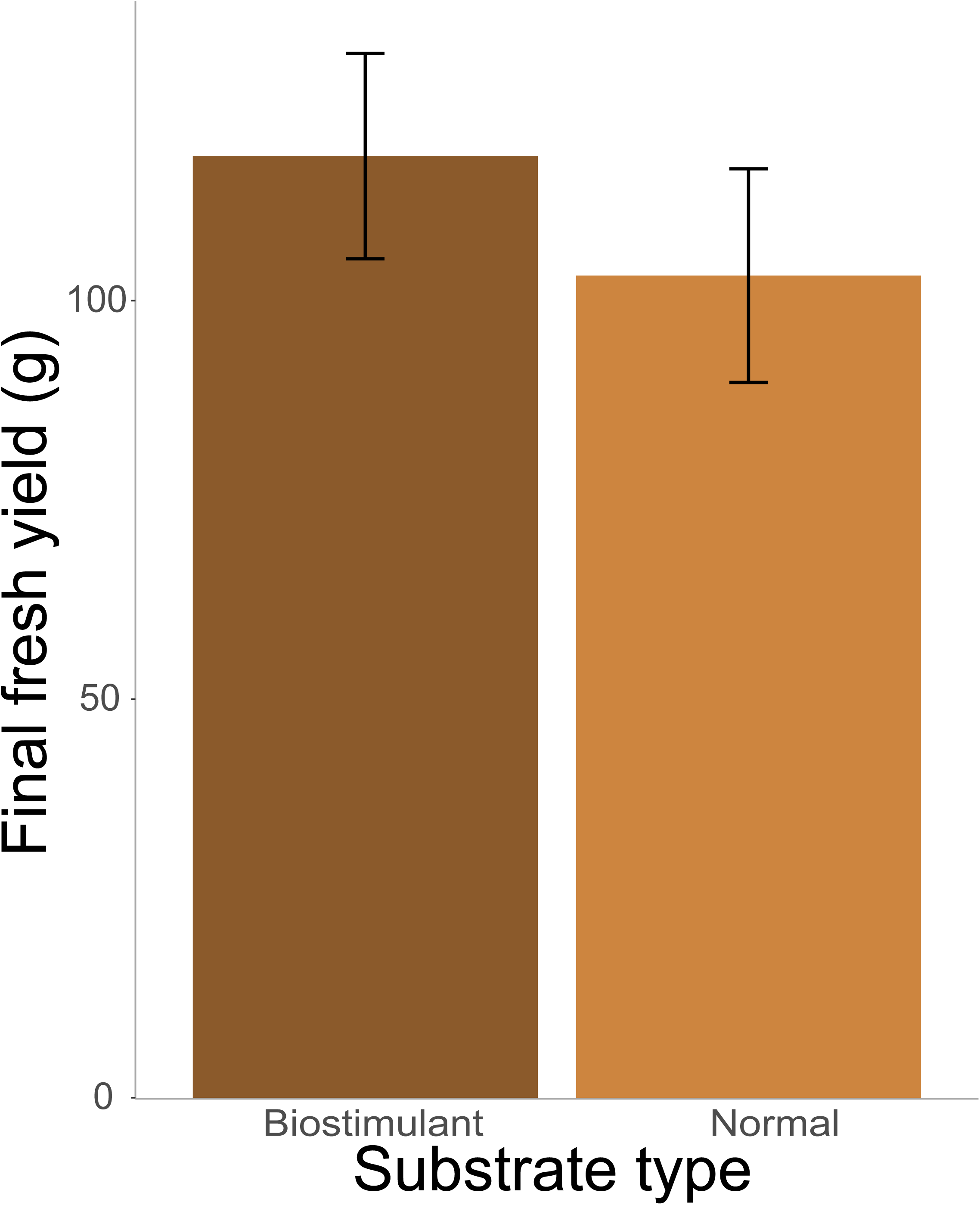
Mean final fresh yields of the cannabis plants after eight weeks according to the substrate type used (normal or with *Bacillus pumilus* biostimulant). Error bars represent 95% confidence intervals.

## DISCUSSION

In this preliminary article, we report the first quantification of yield loss from damage caused by onion thrips for indoor grown cannabis. We estimated yield losses that are higher than those reported for outdoor cannabis growers in California in a survey on all pests (Wilson et al., 2019). Indoor growing environments are particularly favourable for thrips and, thus, likely increase the risk associated with thrips’ outbreaks. A similar study using three initial infestation levels of thrips found that the western flower thrips (*F. occidentalis* (Pergande)) can significantly reduce yields for greenhouse grown cucumbers, as well as the plant’s growth and photosynthesis rates (Hao et al., 2002). A multitude of factors such as the crop nutritional status, its growing condition and the prevalence of pests and diseases influence yields in a crop and thus make describing yields as a function of a precise pest infestation very difficult (Pereira et al., 2017). This is even more true for pests inflicting indirect damage such as thrips (Hao et al., 2002; Pereira et al., 2017). However, as growing conditions are highly controlled in indoor cannabis production, as they were standardized in between our plants, as climatic variations are minimal, and as our plants were all clones equally treated, we believe the differences observed in this study are most likely due to differences in infestations rates.

The onion thrips have a very diversified range of hosts (Diaz-Montano et al., 2011; Nault, Kain, & Wang, 2014; Stuart et al., 2011) and has long been recognized as a greenhouse pest (Morison, 1957). Considering it has been found on hemp (Cranshaw et al., 2019), it is not surprising that indoor grown cannabis can be added to this long list of hosts. Nonetheless, this is a new piece of valuable information for producers. In onions, it can notably impair bulb weight (Ghosheh & Al-Shannag, 2000) and reduce yields, sometime by more than 50% (Diaz-Montano et al., 2011; Fournier, Boivin, & Stewart, 1995), especially since as little as 10 thrips per plant is sufficient to decrease yields by 7% in greenhouses (Kendall & Capinera, 1987). Onion thrips are particularly known to feed on leaves, causing photosynthesis reduction, distorted plant parts, and reduced bulbs size as well as transmitting viruses, such as the Iris yellow spot virus (family Bunyaviridae, genus *Tospovirus*, IYSV) (Diaz-Montano et al., 2011; Gent et al., 2006; Wu et al., 2013). Damage consisting of yellow dot-shaped scars were observed both on the leaves of our experimental plants and on production plants under outbreak pressure (Fig. 3). These injuries can be considered indirect damage and were almost certainly inflicted by the onion thrips. We believe our observations correspond to the “serious foliage damage” reported by Cranshaw et al. (2019) on hemp. Even though we did not investigate damage extent or photosynthetic rate in our cannabis plants, it can be expected that our reduced yields originate from those indirect feeding damage. Similar injuries and scars are known to reduce the photosynthetic ability of leaves in onions (Diaz-Montano et al., 2011). Little information is available about the transmission of viruses to cannabis plants by thrips but McPartland (1996b) mention that viruses can greatly reduce yields in cannabis and that the onion thrips is one of the worst vector of viruses in this crop. Besides yield losses, we also highlight potential decreases in product value through reduced levels of total THC from highly infested plants.

**Fig. 3.**
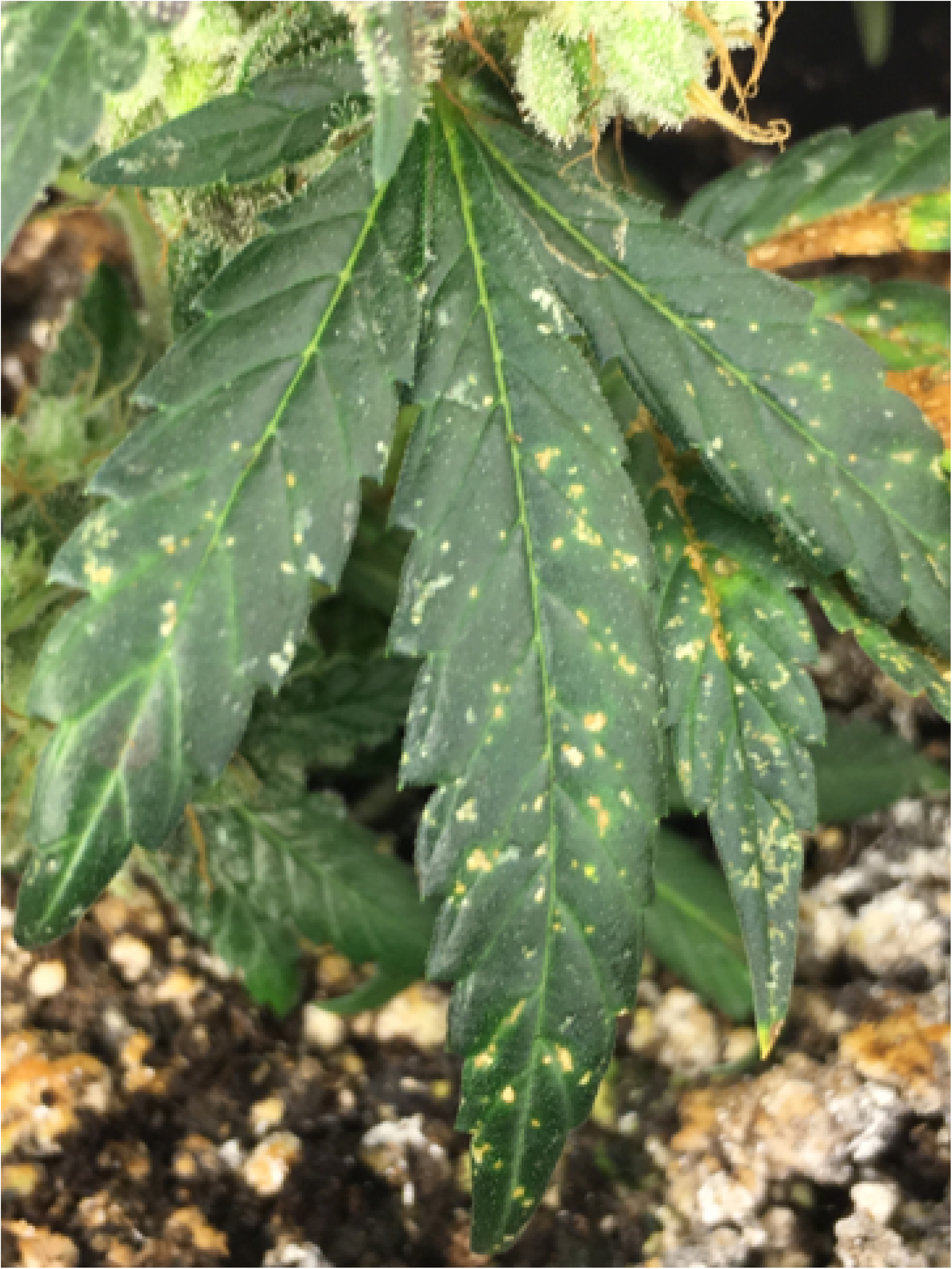
Example of leaf damage observed on the experimental plants and on production plants in the facility.

The use of a growing medium with *B. pumilus* did not improved the plants’ strength. It is consistent with previous experiments with the same enhanced growing medium on the control of the western flower thrips (*F. occidentalis*) (Gravel & Naasz, 2019). However, our results were nearly significant, and we still consider that growing medium enhanced with microbial biostimulants represents a promising avenue for integrated pest management in greenhouse or indoor productions. Their use should be investigated more in indoor cannabis production and for various pests.

In conclusion, we showed that the onion thrips is present in indoor grown cannabis in Canada and that it represents an economic threat. We observed damage caused by thrips feeding on leaves and experimentally found a link between infestation levels and final fresh yields, in addition to a possible reduction of total THC levels under high infestation. This study was preliminary and should motivate further experiments. As chemical means of control are very limited for indoor grown cannabis, we recommend strict monitoring programs for indoor cannabis producers to avoid economic losses. We show that thrips potentially represent a major threat to product quality and yields. In this way, we suggest more research on cannabis pests to identify all pest species in various growing setting, including the range of damage they can cause and their economic thresholds. Only this can subsequently lead to the development of management programs and the development of safe and affordable control methods.

## ACKNOWLEDGMENTS

We thank the team from GreenSeal Cannabis Co., especially Kurt Fisher for providing plant care, and Chris Murray and Peter Reeves for helping build the cages. We are also grateful to Premier Tech for providing the Pro-Mix HP Biostimulant + Mycorrhizae. This research was supported by funding from the NSERC’s Engage Grants for Universities program, no 536612-2018.

